# Long lasting alterations of feeding by adolescent obesity

**DOI:** 10.64898/2026.09.02.748861

**Authors:** Solenne Rougeux, Diptendu Mukherjee, Robert T. West, Kate Z. Peters, Fabien Naneix

## Abstract

The prevalence of obesity is increasing worldwide, especially amongst younger populations. As both brain and body are still under development, numerous studies have shown that adolescence is an especially sensitive period for the impact of external insults including unbalanced diets. To better understand and prevent the vulnerability to obesity and maladaptive food choice, it is crucial to explore the long-term impact of adolescent exposure to obesogenic diets on different feeding behaviours. To address this, we gave mice access to an obesogenic diet with a very high-fat content (vHFD) specifically during adolescence and investigated the long-lasting impact on feeding at adulthood months after a switch to a healthy diet. Using specific feeding tests, we found that both males and females vHFD-exposed mice exhibited an increased intake of both standard food and palatable high-fat foods, without impacting preference for non-caloric food rewards. Importantly, these sex-dependent alterations are not fully correlated with diet-induced weight gain but may support long-term vulnerability to overeating and the development or maintenance of obesity.

## 1. Introduction

Obesity and overweight are among the major health challenges for modern societies. During recent decades the incidence rate of obesity has dramatically increased among younger populations (World Health Organisation, 2025). This is mainly driven by changes in dietary habits, especially the overconsumption of highly palatable and energy-dense foods (e.g. high-fat food, sugary drinks) (Berthoud & Morrison, 2008). Juvenile obesity represents an important risk factor for adult obesity and for the development of obesity-related disease (cardiovascular diseases, diabetes, cancer…) earlier in life (Emerging Risk Factors Collaboration, 2023; Vivier & Tompkins, 2008). To develop better preventive and therapeutic strategies, it is critical to understand the long-term impact of the exposure to obesogenic diets during early life on physiology and behaviour.

Early life periods such as childhood and adolescence are key developmental windows for numerous processes, including the control of feeding and food preferences (Blakemore & Robbins, 2012; Peters & Naneix, 2022; Serrano-Gonzalez et al., 2021; Somerville & Casey, 2010; Spear, 2000). Beyond their impact on body weight and body composition (Guo et al., 2009; Yang et al., 2014), numerous studies have shown that the chronic consumption of obesogenic diets alters numerous processes including food-seeking, food-related decision making and feeding itself (Ferrario et al., 2024; Kenny, 2011; Mazzone et al., 2020; Seabrook et al., 2023; Volkow et al., 2011; Thanarajah et al., 2023). However, such diets can have a greater impact and different effects on brain and behaviour when consumed during specific developmental stages like adolescence (Baker et al., 2017; Lowe et al., 2020; Tsan et al., 2021). Recently we demonstrated that the exposure to high-fat diets during adolescence induces sex-dependent long-lasting alterations of the cognitive control of food-seeking (Mukherjee et al., 2026). The impact on feeding is still unknown, but these alterations suggest long-term vulnerability to obesity and overeating.

Control of feeding has been historically categorised as “homeostatic” eating, e.g. eating to restore energy balance, and “hedonic” eating, e.g. driven by the rewarding, palatable and pleasurable properties of food, seen as separate processes with mostly segregated brain circuits (hypothalamic and brainstem circuits vs dopamine and limbic circuits) (Alcantara et al., 2022; Berridge & Kringelbach, 2015; Berthoud & Morrison, 2008; Ferrario et al., 2024; Rossi & Stuber, 2018). Based on this model, the dysregulation of feeding leading to obesity is often associated with hedonic feeding due to the rewarding properties of obesogenic diets. However, there is increasing evidence that hedonic and homeostatic feeding processes interact and influence each other. For instance, specific physiological states (e.g. starvation, nutrient deficiency) recruit reward brain circuits to drive specific appetite (Fortin & Roitman, 2017; Hsu et al., 2018; Naneix et al., 2020; Onimus et al., 2026; Robinson & Berridge, 2013), while obesogenic diets and obesity itself alters homeostatic feeding processes inducing changes in food preference, energy expenditure and in the consumption of standard foods (Beutler et al., 2020; Ferrario et al., 2024; Mazzone et al., 2020).

Here we investigated the long-lasting impact of a limited exposure to obesogenic diet during adolescence on feeding behaviours at adulthood. Using a range of behavioural tests, we demonstrate that exposure to high-fat diet during adolescence increases different types of feeding in adulthood without impacting the intake or preference for non-caloric palatable solutions. Importantly, we show that that these long-term changes in feeding behaviours are sex-dependent but are not fully explained by diet-induced weight gain.

## 2. Material and methods

### 2.1. Subjects

Male and female C57Bl/6J mice were bred at the Medical Research Facility (University of Aberdeen). At weaning (postnatal day, PND, 21) mice were housed in polycarbonate stock cages (450 x 280 x 130 mm, Techniplast 1292N; 4-7 mice per cage according to their sex) for the duration of diet exposure. At adulthood and outside of behavioural testing, they were group housed in polycarbonate cages (369 x 165 x 132 mm, Techniplast 1145; 1-4 mice per cage). Environmental enrichment was provided by tinted polycarbonate tubes and nesting material. They were maintained in a temperature (19-22°C) and humidity (45-55%) controlled environment with a 12 h light/dark cycle (lights on at 7:00 AM) and with food and water available *ad libitum*, except when otherwise stated. All behavioural experiments took place during the light phase. All procedures were performed in accordance with the Animals (Scientific Procedures) Act 1986 and approved by the local Animal Welfare Ethical Review Body (AWERB; Project License P58595877).

### 2.2. Diets

Mice were randomly assigned to one of the two diet conditions for which they were exposed for 5 weeks between PND28 and PND63 (**Figure 1A**) (Peters & Naneix, 2022; Spear, 2000): **1)** Standard diet (SD group) with only access to standard chow diet (3.6 kcal/g; 69% kcal from carbohydrates, 9% lipids, 22% proteins; CMR, SDS), or **2)** Very High-Fat diet (vHFD group) with access to both standard chow and to 60% fat diet (5.2 kcal/g; 20% kcal from carbohydrate, 60% lipids mainly from lard, 20% proteins; D12492, Research Diets; **Table 1**). vHFD was selected based on our previous work (Mukherjee et al., 2026) and its robust weight gain effect in both sexes (Speakman, 2019). All diets were provided *ad libitum*. Animal’s body weight, water and food intake were recorded weekly. At PND63, all mice were switched to standard chow diet and water only and remained like this for 4 months before behavioural testing (∼PND180). A third diet, High-fat diet (HFD; 4.7 kcal/g; 35% kcal from carbohydrate, 45% lipids mainly from lard, 20% proteins; D12451, Research Diets; **Table 1**) was used for feeding tests (see below).

**Figure 1.**
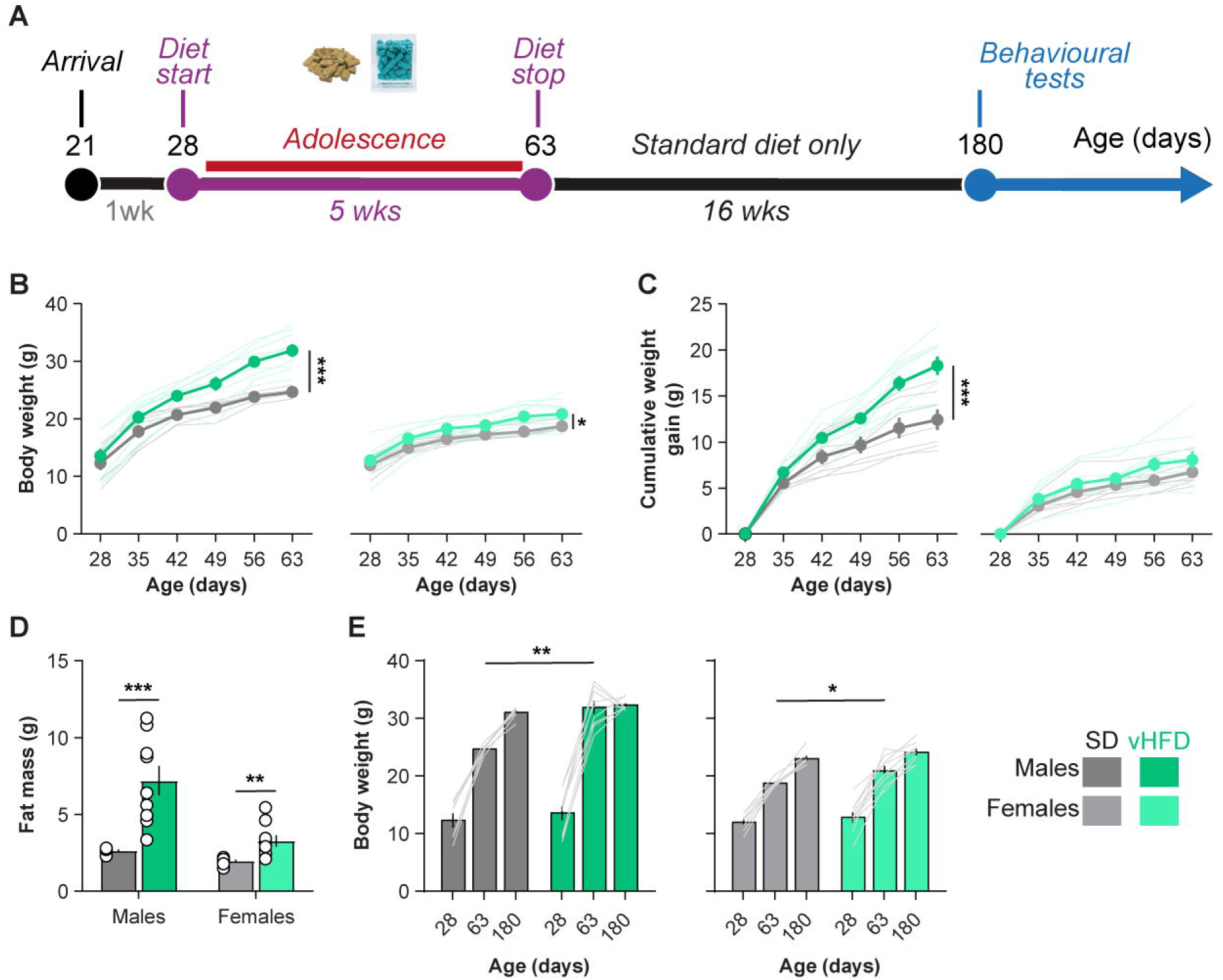
Obesogenic diet during adolescence induces sex-specific impact on body weight. **A.** Schematic representation of experimental design. Mice had continuous access to either standard chow diet only (SD, 9% kcal from fat) or access to both SD and very high-fat diet (vHFD, 60% kcal from fat) during adolescence (postnatal days 28-63). At adulthood, all mice had only access to SD for 4 months before being behavioural testing. **B-D.** vHFD exposure during adolescence increases body weight in both male and female mice (**B-C**) and increases fat mass **(D). E.** Switch to SD normalizes body weight in both males and females before behavioural testing. SD, 7M/8F (black/gray respectively); vHFD, 9M/9F (dark and light green). Data are represented as mean ± SEM (bars or closed circles) with individual values (open circles or thin lines). *, **, *** p < 0.05, 0.01 and 0.001 Diet effect (one-way ANOVA followed by Bonferroni’s post hoc tests or unpaired t-tests). Full statistical reporting is provided in **Supplemental Table 1** Mouse illustration from NIAID NIH BioArt Source (bioart.niaid.nih.gov/bioart/20).

**Table 1.** Macronutrient breakdown of SD, HFD and vHFD.

|  | SD<br>(CMR,<br>SDS) | HFD<br>(D12451,<br>Research<br>Diets) | vHFD<br>(D12492,<br>Research<br>Diets) |
| --- | --- | --- | --- |
|  | 3.6 kcal/g | 4.73 kcal/g | 5.24 kcal/g |
| **<u>Protein</u>** | **22** | **20** | **20** |
| **<u>Carbohydrate</u>** | **69** | **35** | **20** |
| *Starch & fibres* | 65 | 7 | 0 |
| *Sucrose* | 4 | 10 | 7 |
| *Maltodextrin* | 0 | 18 | 13 |
| **<u>Fat</u>** | **9** | **45** | **60** |
| *Lard* | 54 | 39 | 54 |
| *Other* | 6 | 6 | 6 |

### 2.3. Behavioural procedures

All behavioural tests were conducted during the light phase of the light/dark cycle (usually between 9am and 3pm, 5-7 days a week).

#### Feeding tests

Mice were habituated to individual consumption cages (369 x 165 x 132 mm, Techniplast 1145 + cage liner sheet) for 2 days before the start of the feeding tests (1h/day before being replaced in their group housing cages). To prevent any food neophobia, all mice were exposed to the three different diets directly in their home cage 2-3 days before the start of the tests. During each of the following tests, mice were placed in their individual cages with 5-6 g of a specific food in a Petri plastic dish. Food intake measurements were taken after 1, 2, 3 and 6 h. Mice were rehoused in groups at the end of the 6h. Three types of food were tested in the following order: standard chow, very high-fat and high-fat diet (see **Diets** section for nutritional profiles). Each food was tested twice: once with previous *ad libitum* access to food, and once as post-fast refeeding (15 h overnight fasting). All tests were separated by at least 48 h during which they had *ad libitum* access to the chow diet.

#### Saccharin preference tests

For two bottles preference tests, mice were single housed and had access to 75 ml plastic bottles with metal sipper spouts (Classic Pet Products). Spillage was estimated using empty bottles placed in empty cases and intake measures were corrected by this amount (∼0.4 g). Mice were habituated by initially giving them access to two bottles of water for 3-4 days. Mice were then given a series of 48-h 2-bottle saccharin versus water tests at ascending concentrations (0.001, 0.01, 0.1, 1% w/v; Saccharin sodium salt hydrate, S1002, Merck). Position of the two bottles was counterbalanced and swapped daily. Daily intakes were averaged over 2 days at each solution concentration and used to calculate saccharin preference (saccharin intake/total intake x 100).

#### Open-field (OF) and Zero Maze (ZM) tests

Mice were individually placed in a rectangular arena (OF: 40 x 40 x 32 cm, with opaque Perspex walls) or in an elevated zero maze (ZM: diameter 50 cm, elevation 70 cm, with 2 hidden and 2 exposed zones) which they could explore freely for 10 min. Starting position was identical for all mice. The overall path length travelled, the average speed and the time spent in centre versus periphery (OF; central zone 20 x 20 cm), or in exposed zones versus hidden zones (ZM) were recorded using webcams (Logitech C310) coupled with ANY-maze video tracking system. Mazes were cleaned with 70% ethanol between animals.

### 2.4. Experimental design and statistical analysis

Statistical analyses were conducted using GraphPad Prism 11 and SPSS (IBM). Figures were created using GraphPad Prism 11 and Adobe Illustrator. Data are represented as mean ±SEM and individual values unless stated otherwise. Group sizes were estimated based on our previous work. Normality was measured with Shapiro–Wilk test. Based on the robustness of ANOVA to slight non-normality (Knief & Forstmeier, 2021), we opted to use parametric tests for consistency between experiments. Homogeneity of variance was measured with Levene’s test.

Data were analysed by independent Student’s t-tests, or two- or three-way analysis of variance (ANOVA) with or without repeated measures. Greenhouse–Geisser correction was applied when there was unequal variance between conditions for repeated measures. Bonferroni corrected *post hoc* tests for multiple comparisons were performed when appropriate. Pearson’s correlation tests and linear regressions were used to investigate relationships between different body weight and behavioural measurements.

All data were initially compared with Sex as factor. Due to initial differences in diet-induced weight gain and our previous work (Mukherjee et al., 2026), separate analyses for each sex were systematically performed as planned comparisons. Measures of effect size (eta squared η^2^ or partial eta squared η_p_^2^ for ANOVA, Cohen’s d for between-subject contrasts) are stated for each comparison. The alpha risk for the rejection of null hypothesis was set at 0.05. Full statistical reporting is provided in **Supplemental Table 1**.

## 3. Results

### 3.1. Obesogenic diet during adolescence transiently increases body weight

We first measured the impact of exposure to vHFDs during adolescence on body weight and body composition (SD: n = 7 males / 8 females; vHFD; n = 9/9). Both SD male and female mice gained significant weight during adolescence (**Figure 1B**; Males: Week F_(1.77,24.74)_ = 272.9, p < 0.001, η_p_^2^ = 1.0; Females: Week F_(1.61,24.18)_ = 143.5, p < 0.001, η_p_^2^ = 0.9). As previously reported using similar model (Mukherjee et al., 2026), exposure to vHFD during adolescence significantly increased body weight (**Figure 1B**; Males: Diet F_(1,14)_ = 11.2, p = 0.005, η_p_^2^ = 0.4; Week x Diet F_(1.77, 24.74)_ = 10.7, p < 0.001, η_p_^2^ = 0.4 / Females: Diet F_(1,15)_ = 6.6, p = 0.02, η_p_^2^ = 0.3; Week x Diet F = 1.7, p = 0.2, η_p_^2^ = 0.1 / **Figure 1C**; Males t_(14)_ = 3.9, p < 0.001, *d* = 2.0 / Females t_(15)_ = 1.1, p = 0.1, *d* = 0.6). Consistently, vHFD-induced weight gain was associated with increased fat mass (**Figure 1D**; Males t_(14)_ = 4.2, p < 0.001, *d* = 2.1 / Females t_(15)_ = 3.3, p = 0.003, *d* = 1.6). Importantly, higher body weight in vHFD mice, relative to SD mice, was only observed during diet exposure (**Figure 1E**; Start of diet: Males t_(14)_ = 0.7, p = 1.0, *d* = 0.4 / Females t_(15)_ = 0.8, p = 0.4, *d* = 0.4; End of diet: Males t_(14)_ = 5.6, p = 0.002, *d* = 2.8 / Females t_(15)_ = 2.5, p = 0.03, *d* = 1.2) and the difference in body weight was not maintained after mice were switched to SD for 4 months (Males t_(14)_ = 3.9, p = 1.0, *d* = 1.3 / Females t_(15)_ = 1.4, p = 0.2, *d* = 0.7). At adulthood (> PND 70), all mice only had access to SD and were maintained like this for at least 4 months before behavioural testing.

### 3.2. Long-lasting impact of adolescent obesogenic diet on standard diet feeding

At adulthood, we first examined standard feeding through consumption of SD during individual 6h feeding test in mice who were maintained with *ad libitum* access to SD before test day. vHFD exposed male and female mice consumed higher amounts of standard chow compared to controls (**Figure 2A-B**). For males, higher chow intake was observed throughout the whole test, despite only being statistically significant for the final measure (**Figure 2C**; 1h t_(13)_ = 2.0, p = 0.06, *d* = 1.1; 2h t_(13)_ = 1.6, p = 0.1, *d* = 0.8; 3h t_(13)_ = 2.1, p = 0.05, *d* = 1.1; 6h t_(13)_ = 2.5, p = 0.03, *d* = 1.3). Chow intake rate was constant throughout the session but with vHFD males being significantly higher (**Figure 2D**; Block F_(1,13)_ = 0.2, p = 0.7, η_p_^2^ = 0.02; Diet F_(1,13)_ = 6.2, p = 0.03, η_p_^2^ = 0.3; Block x Diet F_(1, 13)_ = 0.2, p = 0.7, η_p_^2^ = 0.02).

**Figure 2.**
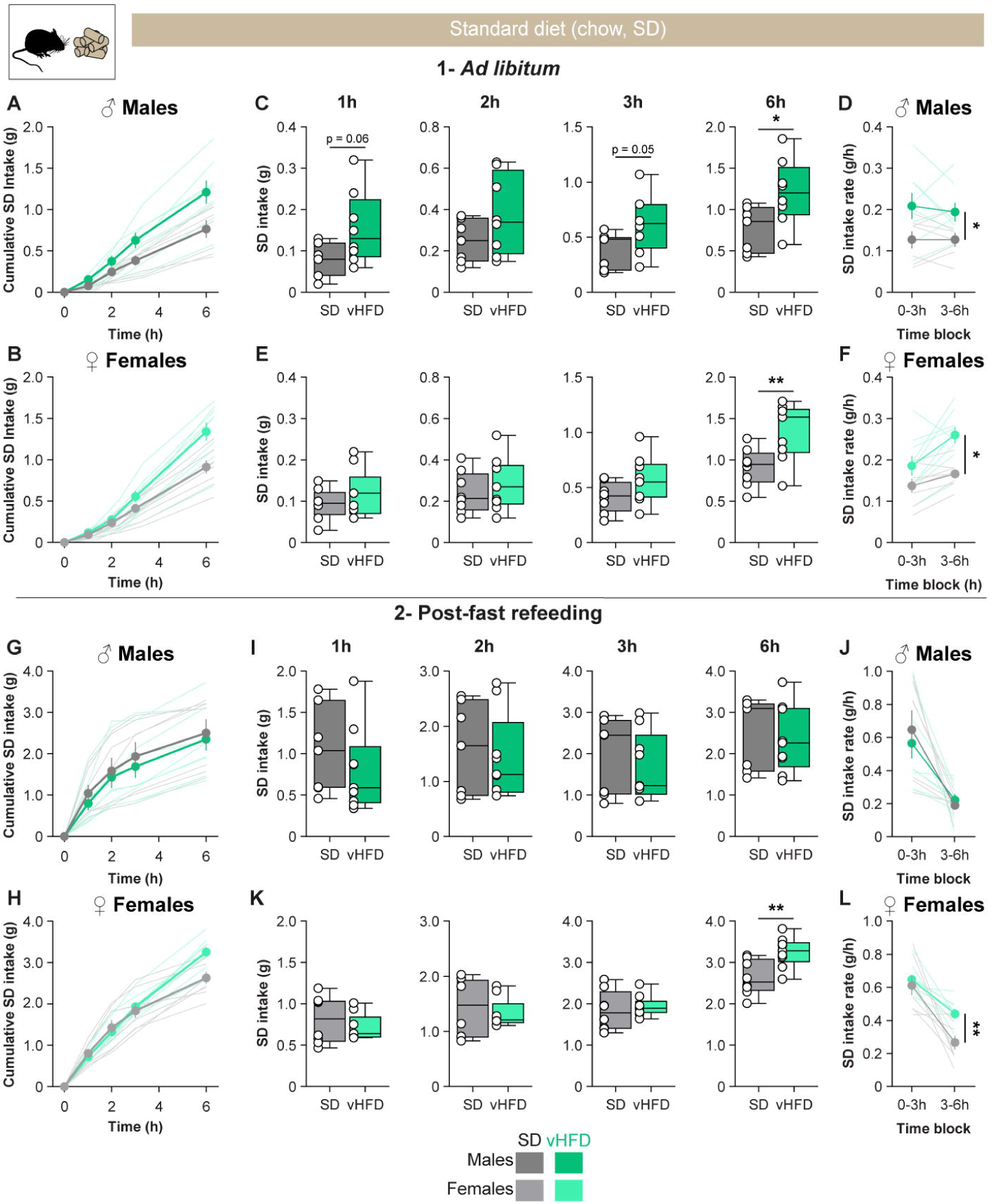
Long-lasting impact of adolescent obesogenic diet on standard diet feeding. **A-B.** Cumulative intake of standard diet (SD, chow) during a 6h feeding test in male (**A**) and female (**B**) mice with prior ad libitum access to food. **C-F.** Cumulative SD intake at 1, 2, 3 and 6h is higher in vHFD-exposed males **(C)** and females **(E),** due to a higher intake rate throughout the test **(D-F). G-H.** Cumulative intake of standard diet (SD, chow) during a 6h feeding test in male (**G**) and female (**H**) mice after overnight fasting. **I-L.** During refeeding, SD and vHFD males ate similar amount of SD at 1, 2, 3 and 6h (**l-J**), while vHFD-exposed females showed higher intake during the second half of the test (**K-L**). SD, 7M/8F (black/gray respectively); vHFD, 8-9M/9F (dark and light green; one vHFD male was excluded during the first test due to an absence of food intake). Data are presented as mean ± SEM (closed circles) or as median and interquartile range (boxplots) with individual values (open circles or thin lines). " p < 0.05 and 0 01 respectively Diet effect (unpaired t-tests or 2-way ANOVAs). Full statistical reporting is provided in **Supplemental Table 1**. Mouse illustration from NIAID NIH BioArt Source (bioart.niaid.nih.gov/bioart/20).

vHFD females also consumed higher cumulative amount of chow compared to their controls (**Figure 2E**; 1h t_(15)_ = 1.1, p = 0.3, *d* = 0.5; 2h t_(15)_ = 0.7, p = 0.5, *d* = 0.3; 3h t_(15)_ = 1.6, p = 0.1, *d* = 0.8, 6h t_(15)_ = 3.0, p = 0.009, *d* = 1.5). Intake rate increases throughout the test but remains higher in vHFD mice (**Figure 2F**; Diet F_(1,15)_ = 251.0, p < 0.001, η_p_^2^ = 0.9; Block F_(1,15)_ = 16.3, p < 0.001, η_p_^2^ = 0.5; Diet x Block F_(1,15)_ = 3.2, p = 0.09, η_p_^2^ = 0.2).

We then tested mice after overnight fasting (post-fast refeeding) to investigate how increased motivational drive affects standard diet feeding. As expected, all mice had higher chow intake after fasting (**Figure 2G-H**). Interestingly, SD and vHFD males exhibited similar refeeding intake during the whole test (**Figure 2I**; 1h t_(14)_ = 0.9, p = 0.4, *d* = 0.5; 2h t_(14)_ = 0.4, p = 0.7, *d* = 0.2; 3h t_(14)_ = 0.6, p = 0.6, *d* = 0.3; 6h t_(14)_ = 0.4, p = 0.7, *d* = 0.2). Both groups have similar intake rates which are decreasing between the first and second half of the test (**Figure 2J**; Diet F_(1,14)_ = 0.1, p = 0.7, η_p_^2^ = 0.6; Block F = 23.7, p < 0.001, η_p_^2^ = 0.6; Block x Diet F_(1, 14)_ = 0.5, p = 0.5, η_p_^2^ = 0.03).

In contrast, vHFD female mice showed higher chow intake during refeeding compared to the SD group (**Figure 2K**; 1h t_(15)_ = 0.8, p = 0.4, *d* = 0.4; 2h t_(15)_ = 0.5, p = 0.6, *d* = 0.2; 3h t_(15)_ = 0.6, p = 0.6, *d* = 0.3; 6h t_(15)_ = 3.3, p = 0.005, *d* = 1.6). As for the males, intake rates progressively decreased during the test but globally remained higher in vHFD females (**Figure 2L**; Diet F_(1,15)_ = 10.9, p = 0.005, η_p_^2^ = 0.4; Block F_(1,15)_ = 35.7, p < 0.001, η_p_^2^ = 0.7, Block x Diet F_(1,15)_ = 0.2, p = 0.1, η_p_^2^ = 0.1).

Taken together, these results demonstrate that adolescent obesity induces long-lasting general alterations of eating for standard diet, while not impairing homeostatic feeding in response to acute energy demands (overnight fasting).

### 3.3. Long-lasting impact of adolescent obesogenic diet on palatable diet feeding

We then evaluated reward-based feeding independent of hunger drive using high-calorie highly palatable foods. Mice with *ad libitum* access to chow diet before test day were first tested with very high-fat food pellets (60% kcal from fat; **Figure 3A-B**). In males, intake amounts were initially similar between SD- and vHFD-exposed groups (**Figure 3C**; 1h t_(14)_ = 0.7, p = 0.5, *d* = 0.4; 2h t_(14)_ = 0.7, p = 0.5, *d* = 0.4; 3h t_(14)_ = 0.7, p = 0.5, *d* = 0.4). However, vHFD-exposed males ate a greater amount of very high-fat food by the end of the test (6h t_(14)_ = 2.5, p = 0.01, *d* = 1.3). This was due to an overall higher food intake rate (**Figure 3D**; Males - Diet F_(1,14)_ = 6.2, p = 0.03, η_p_^2^ = 0.3; Block F_(1,14)_ = 54.5, p < 0.001, η_p_^2^ = 0.8; Block x Diet F_(1, 14)_ = 1.4, p = 0.3, η_p_^2^ = 0.09).

**Figure 3.**
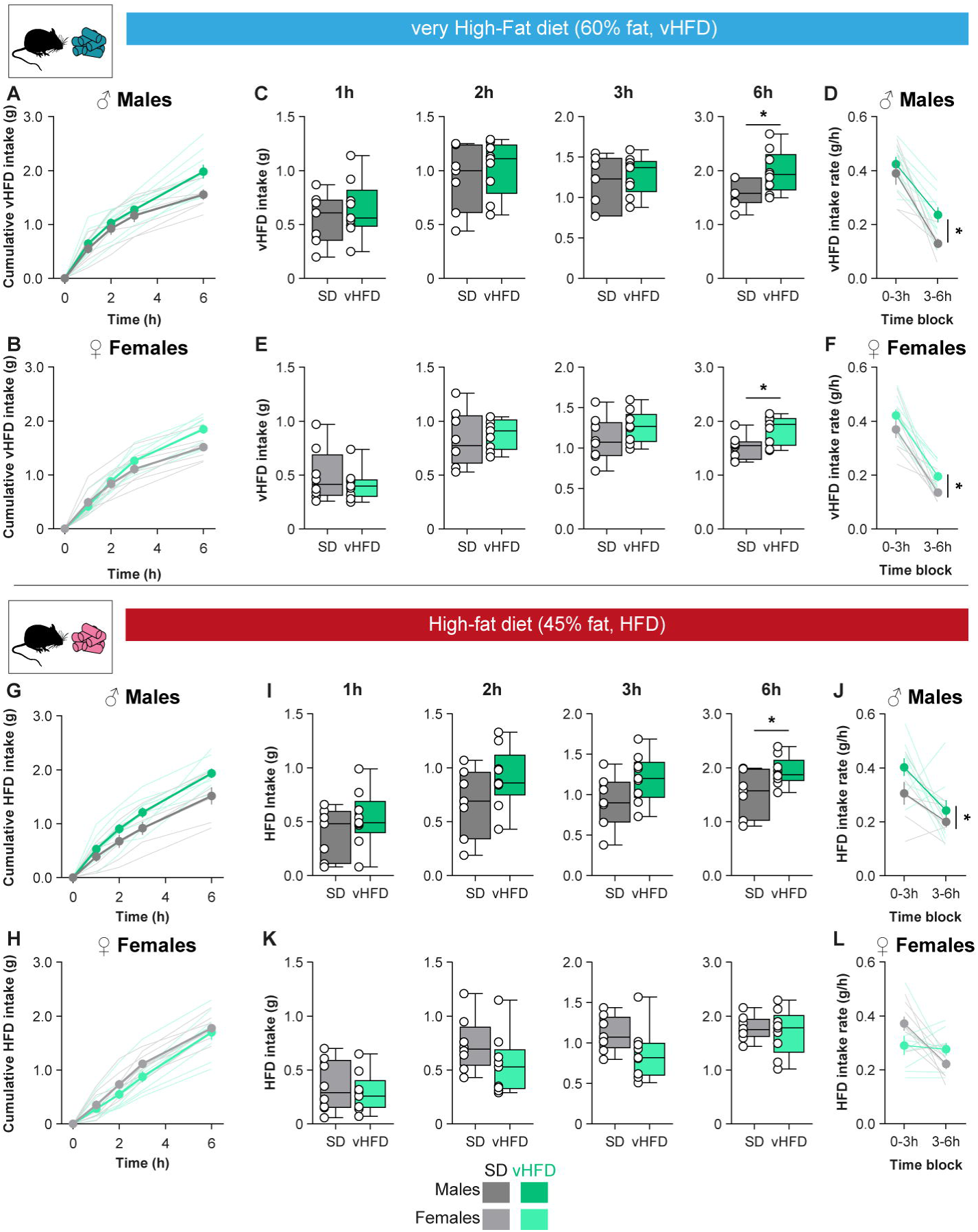
Long-lasting impact of adolescent obesogenic diet on palatable diets feeding. **A-B.** Cumulative intake of very high-fat diet (vHFD) during a 6h feeding test in male (**A**) and female (**B**) mice with prior ad libitum access to chow diet. **C-F.** Cumulative SD intake at 1,2, 3 and 6h is higher in vHFD-exposed males **(C)** and females **(E),** due to a higher intake rate throughout the test **(D-F). G-H.** Cumulative intake of high-fat diet (HFD) during a 6h feeding test in male (**G**) and female (**H**) mice with prior ad libitum access to chow diet. **I-L.** Males (**l-J**) but not females (**K-L**), ate higher amount of HFD, due to higher intake rate throughout the test. SD, 7M/8F (black/gray respectively); vHFD, 9M/9F (dark and light green). Data are presented as mean ± SEM (closed circles) or as median and interquartile range (boxplots) with individual values (open circles or thin lines). * p < 0.05 respectively Diet effect (unpaired t-tests or 2-way ANOVAs). Full statistical reporting is provided in **Supplemental Table 1**. Mouse illustration from NIAID NIH BioArt Source (bioart. niaid.nih.gov/bioart/20).

As the males, female groups showed an initial similar intake (**Figure 3E**; 1h t_(15)_ = 0.8, p = 0.4, *d* = 0.4; 2h t_(15)_ = 0.4, p = 0.7, *d* = 0.2; 3h t_(15)_ = 1.4, p = 0.2, *d* = 0.7), but an overall higher consumption of very high fat food by the end of the test (6h t_(15)_ = 2.8, p = 0.01, *d* = 1.3). Intake rates were overall higher in the vHFD female group (**Figure 3F**; Females - Diet F_(1,15)_ = 7.9, p = 0.01, η_p_^2^ = 0.3; Block F_(1,15)_ = 87.3, p < 0.001, η_p_^2^ = 0.9; Block x Diet F_(1, 15)_ = 0.02, p = 0.9, η_p_^2^ = 0.001).

To control for familiarity with specific diets we used a third diet (high fat food; 45% kcal from fat) which mice had no prior experience with (**Figure 3G-H**). In males, vHFD-exposed mice showed a higher intake of HFD, which is noticeable already during the first three hours (**Figure 3I**; 1h t_(14)_ = 1.1, p = 0.3, *d* = 0.6; 2h t_(14)_ = 1.6, p = 0.2, *d* = 0.8; 3h t_(14)_ = 1.9, p = 0.08, *d* = 0.9; 6h t_(14)_ = 2.4, p = 0.03, *d* = 1.2 / **Figure 3J**; Diet F_(1,14)_ = 5.9, p = 0.03, η_p_^2^ = 0.3; Block F_(1,14)_ = 10.6, p = 0.006, η_p_^2^ = 0.4; Block x Diet F_(1, 14)_ = 0.5, p = 0.5, η_p_^2^ = 0.02), confirming an increase in hedonic feeding.

Interestingly, this pattern was not observed in females where control and vHFD-exposed animals consumed similar amounts of HFD throughout the feeding test (**Figure 3K**; 1h t_(15)_ = 0.6, p = 0.5, *d* = 0.3; 2h t_(15)_ = 1.5, p = 0.2, *d* = 0.7; 3h t_(15)_ = 1.8, p = 0.09, *d* = 0.9; 6h t_(15)_ = 0.5, p = 0.6, *d* = 0.2 / **Figure 3L**; Diet F_(1,15)_ = 0.2, p = 0.7, η_p_^2^ = 0.01; Block F_(1,15)_ = 9.8, p = 0.007, η_p_^2^ = 0.4; Block x Diet F_(1, 15)_ = 6.7, p = 0.02, η_p_^2^ = 0.3; Bonferroni’s *post hoc* tests SD vs vHFD: 0-3h p = 0.09, 3-6h p =0.09).

Taken together, this demonstrates that vHFD mice showed an increase in the consumption of different high-calorie highly palatable foods, which can participate in long-term vulnerability to obesity development.

### 3.4. Adolescent obesogenic diet does not impact intake of non-caloric foods

These findings showed that adolescent exposure to obesogenic diets enhances food intake and the consumption of palatable foods specifically. This raised the question of whether vHFD or HFD consumption during adolescence would also induce a general increase of the consumption of palatable but non-caloric foods. We investigated this using two-bottle preference tests with increasing concentrations of the non-caloric sweetener saccharin (0.001 to 1%; **Figure 4**).

**Figure 4.**
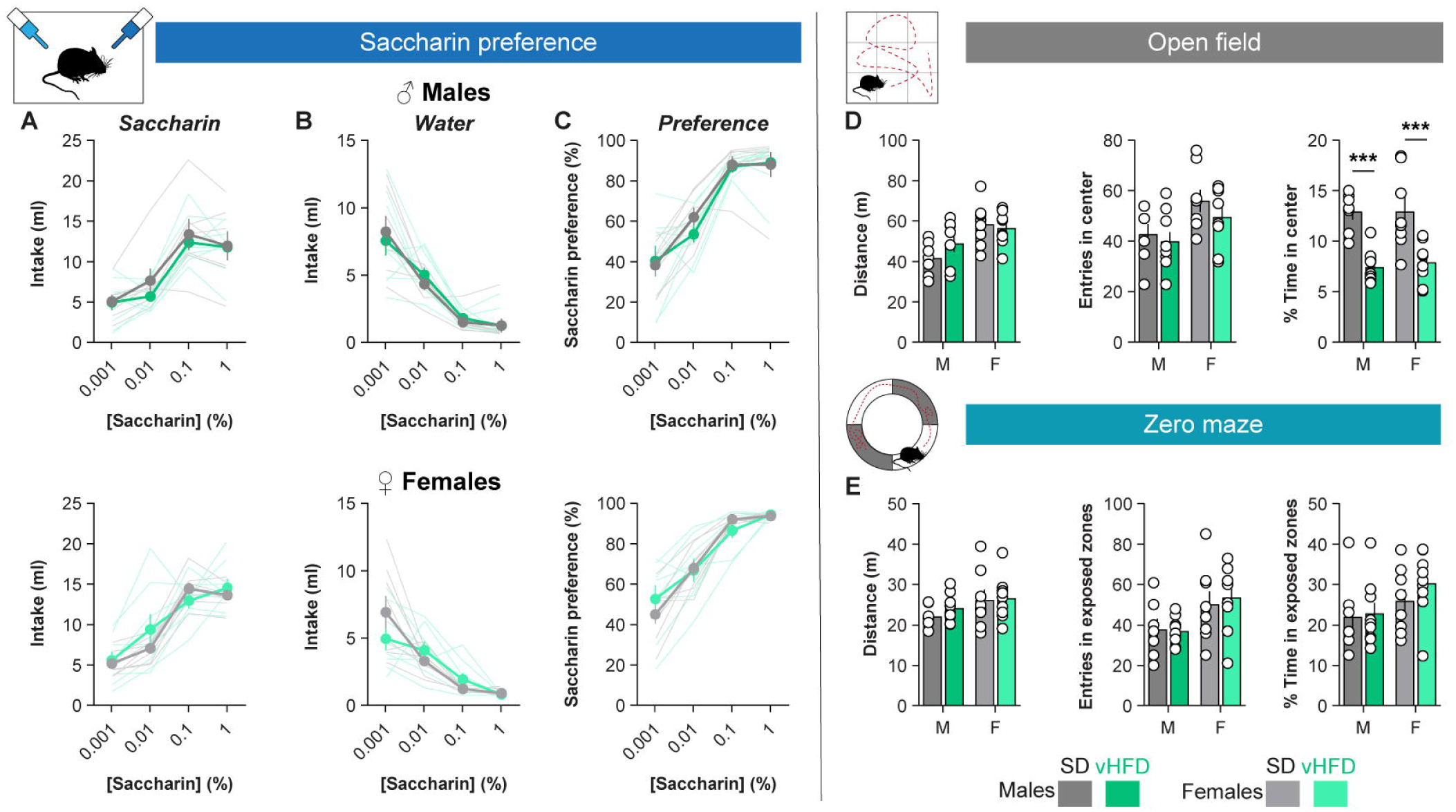
Adolescent obesogenic diet does not impact sweet preference but increases anxiety-associated behaviours. **A-C.** Sweet preference 2-bottles test including saccharin intake (**A**), water intake **(B)** and saccharin preference **(C)** in male (top) and female mice *(bottom).* **D-E.** Anxiety-related behaviours measured in open field **(D)** and zero-maze **(E)** tests including total distance travelled *(left),* total number of entries in centre or exposed zones *(mid)* and percentage of time spent in centre/exposed zones *(right).* SD, 7M/8F (black/gray respectively); vHFD, 9M/8-9F (dark and light green; one vHFD female was excluded during the open field test as it was not exploring). Data are presented as mean ± SEM (closed circles) with individual values (open circles or thin lines). *** p < 0.001 Diet effect (unpaired t-tests). Full statistical reporting is provided in **Supplemental Table 1**. Mouse illustration from NIAID NIH BioArt Source (bioart.niaid.nih.gov/bioart/20).

All male and female mice drank similar volumes throughout the tests with a progressive increase of their saccharin consumption (**Figure 4A**; Males: Diet F_(1, 14)_ = 0.4, p = 0.6, η_p_^2^ = 0.03; Concentration F_(1.69, 23.59)_ = 41.1, p < 0.001, η_p_^2^ = 0.8; Diet x Concentration F = 0.5, p = 0.6, η_p_^2^ = 0.04 / Females: Diet F_(1, 15)_ = 0.005, p = 1.0, η_p_^2^ = 0; Concentration F_(3, 45)_ = 56.5, p < 0.001, η_p_^2^ = 0.8; Diet x Concentration F_(3, 45)_ = 2.5, p = 0.07, η_p_^2^ = 0.1) and parallel decrease of water intake (**Figure 4B**; Males: Diet F_(1, 14)_ = 0.01, p = 0.9 η_p_^2^ = 0.001; Concentration F_(1.36, 19.06)_ = 45.8, p < 0.001, η_p_^2^ = 0.8; Diet x Concentration F_(1.36, 19.06)_ = 0.4, p = 0.6, η_p_^2^ = 0.2 / Females: Diet F_(1, 15)_ = 0.2, p = 0.7, η_p_^2^ = 0.01; Concentration F_(1.74, 26.15)_ = 30.4, p < 0.001, η_p_^2^ = 0.7; Diet x Concentration F_(1.74, 26.15)_ = 2.8, p = 0.09, η_p_^2^ = 0.2). Taken together, this resulted in a concentration-dependent increase in saccharin preference (**Figure 4A**; Males: Concentration, F_(1.54, 21.58)_ = 57.2, p < 0.001, η_p_^2^ = 0.8 / Females: Concentration F_(2.13, 31.88)_ = 71.1, p < 0.001, η_p_^2^ = 0.8) similar in all groups (Males: Diet F_(1, 14)_ = 0.2, p = 0.7, η_p_^2^ = 0.01; Diet x Concentration F_(1.54, 21.58)_ = 0.6, p = 0.5, η_p_^2^ = 0.04 / Females: Diet F_(1, 15)_ = 0.4, p = 0.6, η_p_^2^ = 0.02; Diet x Concentration F_(2.13, 31.88)_ = 1.3, p = 0.3, η_p_^2^ = 0.08). Thus, adolescent exposure to vHFD did not induce long-lasting disruptions in sensitivity to sweet taste and the consumption of non-caloric palatable foods.

### 3.5. Long-term impact of adolescent obesogenic diet on anxiety-related behaviours

Finally, the consumption of obesogenic diets is known to promote anxiety-like behaviours (Tsan et al., 2021). Thus, we investigated the response of the vHFD-exposed mice to two tests classically used to explore such behaviours: the open field (OF) and the elevated zero-maze (ZM; **Figure 4D-E**). In both tests, SD- and vHFD-exposed mice exhibited similar general locomotor activity (OF: Males t_(14)_ = 1.4, p = 0.2, *d* = 0.7; Females t_(14)_ = 0.4, p = 0.7, *d* = 0.2 / ZM: Males t_(14)_ = 1.3, p = 0.2, *d* = 0.6; Females t_(15)_ = 0.1, p = 0.9, *d* = 0.06) and a similar number of entries in the centre (OF: Males t_(14)_ = 0.5, p = 0.6, *d* = 0.3; Females t_(14)_ = 1.6, p = 0.1, *d* = 0.8) and in exposed areas (ZM: Males t_(14)_ = 0.2, p = 0.9, *d* = 0.08; Females t_(15)_ = 0.4, p = 0.7, *d* = 0.2). During the OF test vHFD-exposed mice spent less time in the centre (Males t_(14)_ = 6.2, p < 0.001, *d* = 3.1; Females t_(14)_ = 3.3, p = 0.006, *d* = 1.6). However, similar anxiety-related behaviour was not observed during the ZM test (Males t_(14)_ = 0.2, p = 0.9, *d* = 0.1; Females t_(15)_ = 1.1, p = 0.3, *d* = 0.5). Thus, while results from the OF test might point toward a long-lasting increase in anxiety-related behaviours, results from the ZM test prevent definitive conclusion.

### 3.6. Sex-dependent relationships between adolescent obesity and long-lasting alterations of homeostatic and hedonic feeding

We then explored the relationships between diet-related observable effects (body weight related measures, fat mass), feeding (standard and palatable foods, saccharin preference) and anxiety-related behaviours in males and females separately (**Figure 5 A-B**).

**Figure 5.**
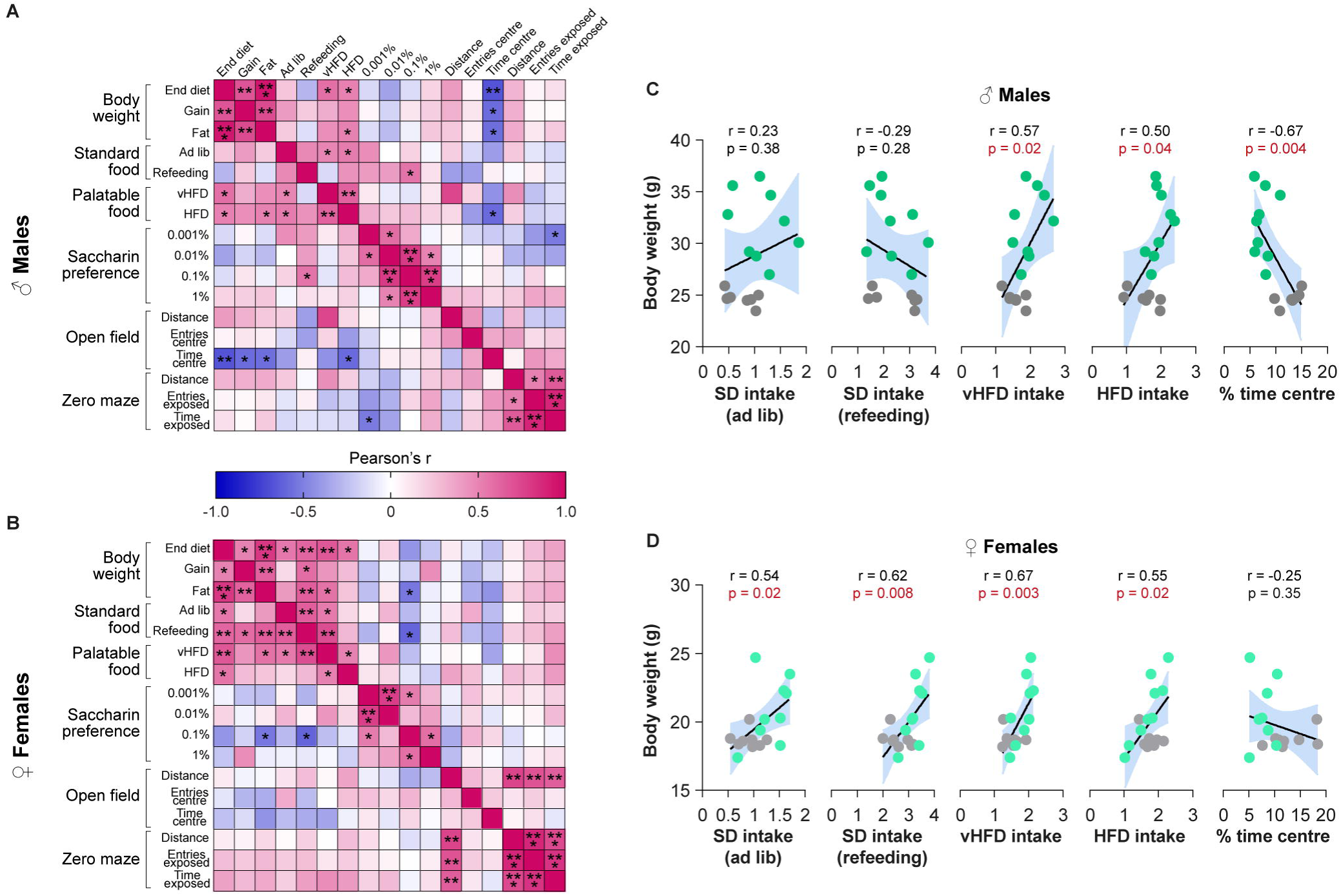
Sex-dependent relationships between adolescent obesity and long-lasting alterations of feeding, anhedonia and anxiety-related behaviours. **A-B.** Correlation matrices (as Pearson’s r heatmap) between body weight measures, feeding, saccharin preference and anxiety-related measures for males (fop) and females *(bottom).* **C-D.** Linear regression between mice’ body weight at the end of diet exposure and standard diet feeding (SD intake ad libitum and during refeeding), palatable diets feeding (vHFD and HFD intake), or time spent exploring the centre of the open field. SD (black/gray respectively); vHFD (dark and light green. *, **, *** p < 0.05, 0.01 and 0.001 respectively (Pearson correlation coefficient).

Pearson correlation matrices revealed positive associations between all body weight related variables in both males (r = [0.68; 0.93], all p < 0.01) and females (r = [0.50; 0.80], all p < 0.05). We then used linear regression to predict different behavioural measures based on end of diet body weights. In males, there were significant associations between body weight at the end of diet exposure and feeding of palatable foods (vHFD intake F_(1,14)_ = 6.8, p = 0.02, r = 0.57; HFD intake F_(1,14)_ = 4.7, p = 0.04, r = 0.5) and with the time spent in the centre of the open field (F_(1,14)_ = 11.6, p = 0.004, r = −0.67). However, males’ body weight was not predictive of standard diet intake (all F < 1.3, all p > 0.2, r = [-0.28; 0.23]). In contrast, females’ body weight at the end of diet exposure was predictive of both palatable (vHFD F_(1,15)_ = 12.0, p = 0.003, r = 0.67; HFD F_(1,15)_ = 6.6, p = 0.02, r = 0.55) and standard foods intake (SD *ad libitum* F_(1,15)_ = 6.3, p = 0.02, r = 0.54; SD refeeding F_(1,15)_ = 9.2, p = 0.008, r = 0.62). However, there was no significant association with the time spent in the centre of the open field (F_(1,14)_ = 0.9, p = 0.3, r = 0.3). In both males and females, saccharin preference was never significantly predicted by body weight (all F < 3.1, all p > 0.09, r = [-0.42; 0.18]).

## 4. Discussion

In the present study, we reveal in a mouse model that an obesogenic diet during adolescence induces long-lasting alterations in different forms of feeding at adulthood. This is observed in both males and females despite a transient increase in body weight limited to the diet exposure period. However, intake and sensitivity to non-caloric palatable foods is not impacted by adolescent diet. Taken together this represents a potential vulnerability mechanism to overeating and obesity.

As we previously showed, the increase in body weight and fat mass levels induced by vHFD is sex-dependent (Mukherjee et al., 2026). Consistent with the literature, males were more sensitive to the obesogenic effects of vHFD showing a clear increase of body weight (+148% from PND28 vs +114% for SD) and accumulation of fat mass compared to control. In contrast, females only showed a modest effect (+69% from PND28 vs +60% for SD) (Frias et al., 2001; Macotela et al., 2009; Yang et al., 2014). It is important to note that in our model, vHFD-exposed mice had concurrent and continuous access to both standard (chow) and very high-fat diets during adolescence. We previously reported that in this model vHFD mice adjusted their intake to maintain a similar calorie intake to control animals, but overwhelmingly preferred vHFD as their main source of energy (Mukherjee et al., 2026). This highlights the complex interactions between energy balance, diet nutrient content, sex and how this impacts body weight and cognition (Hu et al., 2018; Noble & Kanoski, 2016; Valladolid-Acebes et al., 2011).

Body weight differences between diet groups did not persist once the diet was removed and mice were tested months after. Studies using healthy dietary interventions after access to obesogenic diets do not report consistent effects on behaviour (Boitard et al., 2016; Hayes et al., 2024; Mazzone et al., 2020; Tran & Westbrook, 2017; Tsan et al., 2022) or metabolism (Guo et al., 2009; Parekh et al., 1998). Here, we did not measure fat mass levels at the time of testing so we cannot fully exclude that vHFD-exposed animals still maintain higher levels of fat mass compared to controls (Guo et al., 2009). However, we previously demonstrated that other measures of metabolic health such as alterations in the response to glucose tolerance test were reversed 2 months after the end of diet exposure (Mukherjee et al., 2026). This strongly suggests that the observed behavioural differences cannot be attributed to ongoing obesity.

The main aim of this study was to investigate the impact of the consumption of obesogenic diets selectively during adolescence on future feeding behaviours. We first showed that both male- and female-exposed mice consumed more standard food (chow) when they were not in a food restricted or food deprived state. This clearly emerged after several hours of intake (6h total test duration), but was already noticeable early on, especially in males. Accordingly, both vHFD-exposed male and female mice showed a higher food intake rate throughout the test. This result strongly contrasts with the devaluation and associated decreased intake of less palatable foods observed during chronic exposure to high-fat foods or shortly after dietary switch to chow (Mazzone et al., 2020). However, this raises the possibility that long-term withdrawal from vHFD sensitises feeding responses including to chow diet (Altherr et al., 2021).As previously stated, we cannot however exclude that these differences are not supported by higher levels of fat mass or energy imbalance in vHFD-exposed mice despite similar body weight after 4 months of diet switch (Guo et al., 2009; Parekh et al., 1998).

During refeeding after overnight fasting, all male mice ate similar total amounts of standard chow food while vHFD-exposed females showed higher intake than their controls. A more detailed analysis showed that vHFD-exposed females showed a similar initial intake (0-3h) but maintained a higher intake rate during the second half of the test (3-6h), resulting ultimately in a higher total intake. In contrast to the feeding test performed with prior *ad libitum* access to food, overnight fasting induced a strong energy deficit and a stronger homeostatic drive to eat. Thus, our results suggest that homeostatic feeding in response to an acute energy deficit is not altered by vHFD adolescent exposure. However, the sustained increase in basal SD intake in energy replete conditions in these animals suggest alterations of other eating processes including changes in meal patterns (initiation, duration, termination), reduced satiation or increased appetition (Alcantara et al., 2022; Rathod & Di Fulvio, 2021; Sclafani, 2013a), which may lead to long-term hyperphagia and weight gain (Heyward & Rosen, 2026). Unfortunately manual weighing of food consumption as used in the present study precludes such in-depth analyses (Ali & Kravitz, 2018). Future studies using automated systems, home cage monitoring or lickometer devices combined with sophisticated video tracking approaches (Ali & Kravitz, 2018; Jhuang et al., 2010; Matikainen-Ankney et al., 2021; Naneix et al., 2020; Taghipourbibalan & McCutcheon, 2026b, 2026a) could explore long-term alterations of mechanism governing ingestive behaviours. This could also reveal potential sex differences as our correlational analyses showed that standard food intake is strongly correlated with diet-related body weight measures only in females.

Using highly palatable foods in food replete conditions, we also showed an increased consumption of food rewards in both males and females (Carlin et al., 2016), which was strongly correlated with weight and weight gain at the end of diet exposure. As with the standard chow, this effect only appears during the second half of the test, suggesting that initial feeding is similar between groups. The sustained feeding rate during the second part of the test suggests more changes in meal initiation and termination and post ingestive processes (Naneix et al., 2020; Smith, 2000). A possible caveat in our experiment was the use of the same food (vHFD) for the adolescent exposure and the palatable food for the feeding tests. However, similar results were observed using another palatable food (HFD) which mice had no prior exposure to, at least in males consistent with sex-specific hyperphagia previously reported for certain palatable foods (Maric et al., 2022). This demonstrates that the increased feeding of palatable foods in vHFD-exposed animals is neither the result from vHFD animals’ previous experience with vHFD nor control animals’ neophobic response to new foods. This result strikingly contrasts to what was observed in animals under obesogenic diets or in people living with obesity showing also an attenuated response to palatable foods and alterations of reward-related behaviours (Arcego et al., 2020; Berland et al., 2021; Íbias et al., 2016; Kenny, 2011; Thanarajah et al., 2023; Volkow et al., 2011) and may support long-term vulnerability to unhealthy dietary habits relapse (van Baak & Mariman, 2023).

Importantly, vHFD-exposed males and females showed similar concentration-dependent consumption and preference for the non-caloric sweetener saccharin. Considering that vHFD also contains higher levels of sucrose compared to SD (**Table 1**), this indicates that vHFD adolescent exposure did not induce long-term changes in sensitivity to sweet tastes or in hedonic perception of sweet rewards. However, it does not rule out other alterations in taste and reward processing including response to dietary lipids (Berland et al., 2020, 2021) and the integration of post-ingestive nutrient sensing (de Araujo et al., 2020; McCutcheon, 2015; McDougle et al., 2024; Sclafani, 2013b; Thanarajah et al., 2019). Such alterations could explain the dysregulation of feeding for both palatable and standard foods seen after several hours of testing. Multiple evidence now points at the ability of the brain reward system to integrate ingestive and post-ingestive signals through different timescales to guide food- and drink-seeking behaviours (de Araujo et al., 2020; Grove et al., 2022; Zhu et al., 2025; Zimmerman & Knight, 2020). These circuits are classically impacted by obesogenic diets and palatable foods including during adolescence (Peters & Naneix, 2022). Thus, it is possible that the adolescent exposure to obesogenic diets induced long-lasting alterations of these integration processes supporting the dysregulation of feeding, food-seeking and food choice behaviours (Thanarajah et al., 2019; van Galen et al., 2023).

Response to stress is a potential factor driving obesity and unhealthy food choices. Exposure to obesogenic diets, especially during early-life, is usually associated with increased anxiety-related behaviours and dysregulation of glucocorticoid hormones later in life (Tsan et al., 2021). Here, we report conflicting results on anxiety-related behaviours using both OF and ZM tests. vHFD-exposed mice spent less time in the centre of the OF suggesting an increase in anxiety-like behaviour. However, similar behaviour was not observed looking at the time spent in the exposed arms of the ZM. Our correlation analyses also showed that measures in both tests are not strongly correlated. The construct validity and the consistency of both OF and ZM tests to measure anxiety-like behaviours is debated (Ennaceur, 2014). Importantly, all mice travelled similar distance in both tests, demonstrating no deficit in locomotor activity in either of the tests. While we cannot totally exclude diet-induced long-term increase in anxiety-related behaviours, decrease in time spent in the centre of the OF can also be interpreted as a deficit in exploratory behaviours despite similar numbers of entries. Future investigations or more in-depth behavioural measurements may give a more precise answer but are beyond the scope of the current study.

In conclusion, we reveal that the chronic consumption of energy-rich obesogenic diets during adolescence leads to long-lasting dysregulation of feeding processes, affecting both homeostatic and hedonic feeding. While most of the impairments reported here were similar in both males and females, subtle differences between sexes highlight the importance of sex as a critical factor for nutrition studies. Taken together with our previous results focusing on the flexible control of decision-making (Mukherjee et al., 2026), this may support a long-term vulnerability to maladaptive food choice and feeding patterns and lead to the maintenance of obesity-promoting behaviours.

## 5. Data availability

Upon publication, all data analysed in this paper will be available on Figshare https://doi.org/10.6084/m9.figshare.33395608

## 6. Acknowledgements

The authors would like to acknowledge the help and support from the staff of the Medical Research Unit, University of Aberdeen for technical support and the care of experimental animals. The authors wish to thank Prof James McCutcheon for helpful discussion and comments on this manuscript.

## 7. Author contributions

SL: Investigation, Writing – Review & Editing; DM: Investigation, Writing – Review & Editing; RTW: Investigation, Validation, Writing – Review & Editing; KZP: Conceptualization, Formal Analysis, Methodology, Visualization, Writing – Review & Editing; FN: Conceptualization, Data Curation, Formal Analysis, Funding Acquisition, Investigation, Methodology, Project Administration, Supervision, Visualization; Writing – Original Draft Preparation, Writing – Review & Editing.

All authors approved the final version of the manuscript.

