## Supplemental Table 1 for "Long lasting alterations of feeding by adolescent obesity"

### Supplemental Table 1: Full statistical report for Figures 1-4

| Figure 1 | | | | | | | | | | | | | |
|---|---|---|---|---|---|---|---|---|---|---|---|---|---|
| **Figure** | **Data** | **Test** | **Factor** | **F-value or t-value** | **p-value** | **Test** | **Factor** | **F-value or t-value** | **p-value** | **Effect size** | **Pairwise comparisons** | | **n (M/F)** |
| **1B** | **Body weight** | 3-way ANOVA | Diet | F(1,29) = 18.0 | p < 0.001 | 2-way ANOVA (Males) | Diet | F(1,14) = 11.2 | p = 0.005 | $\eta_p^2 = 0.4$ | *Post hoc* Bonferroni (Diet) | p < 0.05 from PND42 | SD n = 7/8<br>vHFD n = 9/9 |
| | | | Sex | F(1,29) = 55.4 | p < 0.001 | | Age | F(1.77,24.74) = 272.9 | p < 0.001 | $\eta_p^2 = 1.0$ | | | |
| | | | Age | F(1.76,51.05) = 425.6 | p < 0.001 | | Age x Diet | F(1.77, 24.74) = 10.7 | p < 0.001 | $\eta_p^2 = 0.4$ | | | |
| | | | Age x Sex | F(1.76,51.05) = 54.0 | p < 0.001 | | | | | | | | |
| | | | Age x Diet | F(1.76,51.05) = 11.7 | p < 0.001 | 2-way ANOVA (Females) | Diet | F(1,15) = 6.6 | p = 0.02 | $\eta_p^2 = 0.3$ | *Post hoc* Bonferroni (Diet) | PND42: p = 0.06<br>PND56: p = 0.04<br>All other p > 0.1 | |
| | | | Diet x Sex | F(1,29) = 2.8 | p = 0.1 | | Age | F(1.61,24.18) = 143.5 | p < 0.001 | $\eta_p^2 = 0.9$ | | | |
| | | | Age x Sex x Diet | F(1.76,51.05) = 4.6 | p = 0.02 | | Age x Diet | F(1.61,24.18) = 1.7 | p = 0.2 | $\eta_p^2 = 0.1$ | | | |
| **1C** | **Body weight gain** | 2-way ANOVA (Total weigh gain) | Diet | F(1,29) = 14.6 | p < 0.001 | Unpaired t-test (Males) | | t(14) = 3.9 | p < 0.001 | d = 2.0 | - | - | |
| | | | Sex | F(1,29) = 71.0 | p < 0.001 | | | | | | | | |
| | | | Diet x Sex | F(1,29) = 5.9 | p = 0.02 | Unpaired t-test (Females) | | t(15) = 1.1 | p = 0.1 | d = 0.6 | - | - | |
| **1D** | **Fat mass** | 2-way ANOVA | Diet | F(1,29) = 26.9 | p < 0.001 | Unpaired t-test (Males) | | t(14) = 4.2 | p < 0.001 | d = 2.1 | - | - | |
| | | | Sex | F(1,29) = 16.6 | p < 0.001 | | | | | | | | |
| | | | Diet x Sex | F(1,29) = 8.3 | p = 0.007 | Unpaired t-test (Females) | | t(15) = 3.3 | p = 0.003 | d = 1.6 | - | - | |
| **1E** | **Body weight Start diet** | 2-way ANOVA | Diet | F(1,29) = 1.1 | p = 0.3 | Unpaired t-test (Males) | | t(14) = 0.7 | p = 1.0 | d = 0.4 | - | - | |
| | | | Sex | F(1,29) = 0.3 | p = 0.6 | | | | | | | | |
| | | | Diet x Sex | F(1,29) = 0.04 | p = 0.8 | Unpaired t-test (Females) | | t(15) = 0.8 | p = 0.4 | d = 0.4 | - | - | |
| | **Body weight End diet** | 2-way ANOVA | Diet | F(1,29) = 36.8 | p < 0.001 | Unpaired t-test (Males) | | t(14) = 5.6 | p = 0.002 | d = 2.8 | - | - | |
| | | | Sex | F(1,29) = 121.1 | p < 0.001 | | | | | | | | |
| | | | Diet x Sex | F(1,29) = 10.5 | p = 0.003 | Unpaired t-test (Females) | | t(15) = 2.5 | p = 0.03 | d = 1.2 | - | - | |
| | **Body weight Testing** | 2-way ANOVA | Diet | F(1,29) = 6.2 | p = 0.02 | Unpaired t-test (Males) | | t(14) = 3.9 | p = 1.0 | d = 1.3 | - | - | |
| | | | Sex | F(1,29) = 314.7 | p < 0.001 | | | | | | | | |
| | | | Diet x Sex | F(1,29) = 0.04 | p = 0.8 | Unpaired t-test (Females) | | t(15) = 1.4 | p = 0.2 | d = 0.7 | - | - | |

### Figure 2

| Figure | Data | Test | Factor | F-value or t-value | p-value | Test | Factor | F-value or t-value | p-value | Effect size | Pairwise comparisons | | n (M/F) |
|---|---|---|---|---|---|---|---|---|---|---|---|---|---|
| 2C&E | SD Intake (1h) | 2-way ANOVA | Diet | F(1,28) = 5.6 | p = 0.03 | Unpaired t-test (Males) | | t(13) = 2.0 | p = 0.06 | d = 1.1 | - | - | SD n = 7/8<br>vHFD n = 8/9 |
| | | | Sex | F(1,28) = 0.3 | p = 0.6 | | | | | | | | |
| | | | Diet x Sex | F(1,28) = 1.4 | p = 0.3 | Unpaired t-test (Females) | | t(15) = 1.1 | p = 0.3 | d = 0.5 | - | - | |
| | SD Intake (2h) | 2-way ANOVA | Diet | F(1,28) = 2.9 | p = 0.1 | Unpaired t-test (Males) | | t(13) = 1.6 | p = 0.1 | d = 0.8 | - | - | |
| | | | Sex | F(1,28) = 1.1 | p = 0.3 | | | | | | | | |
| | | | Diet x Sex | F(1,28) = 0.9 | p = 0.4 | Unpaired t-test (Females) | | t(15) = 0.7 | p = 0.5 | d = 0.3 | - | - | |
| | SD Intake (3h) | 2-way ANOVA | Diet | F(1,28) = 7.4 | p = 0.01 | Unpaired t-test (Males) | | t(13) = 2.1 | p = 0.05 | d = 1.1 | - | - | |
| | | | Sex | F(1,28) = 0.06 | p = 0.8 | | | | | | | | |
| | | | Diet x Sex | F(1,28) = 0.5 | p = 0.5 | Unpaired t-test (Females) | | t(15) = 1.6 | p = 0.1 | d = 0.8 | - | - | |
| | SD Intake (6h) | 2-way ANOVA | Diet | F(1,28) = 14.8 | p < 0.001 | Unpaired t-test (Males) | | t(13) = 2.5 | p = 0.03 | d = 1.3 | - | - | |
| | | | Sex | F(1,28) = 1.5 | p = 0.2 | | | | | | | | |
| | | | Diet x Sex | F(1,28) = 0.006 | p = 0.9 | Unpaired t-test (Females) | | t(15) = 3.0 | p = 0.009 | d = 1.5 | - | - | |
| 2D&F | SD intake rate | 3-way ANOVA | Diet | F(1,28) = 14.9 | p < 0.001 | 2-way ANOVA (Males) | Diet | F(1,13) = 6.2 | p = 0.03 | $\eta_p^2 = 0.3$ | - | - | |
| | | | Sex | F(1,28) = 1.5 | p = 0.2 | | Block | F(1,13) = 0.2 | p = 0.7 | $\eta_p^2 = 0.02$ | | | |
| | | | Block | F(1,28) = 4.6 | p = 0.04 | | Block x Diet | F(1, 13) = 0.2 | p = 0.7 | $\eta_p^2 = 0.02$ | | | |
| | | | Block x Sex | F(1,28) = 8.1 | p = 0.008 | | | | | | | | |
| | | | Block x Diet | F(1,28) = 0.5 | p = 0.5 | 2-way ANOVA (Females) | Diet | F(1,15) = 251.0 | p < 0.001 | $\eta_p^2 = 0.9$ | - | - | |
| | | | Diet x Sex | F(1,28) = 0.006 | p = 0.9 | | Block | F(1,15) = 16.3 | p < 0.001 | $\eta_p^2 = 0.5$ | | | |
| | | | Block x Sex x Diet | F(1,28) = 2.2 | p = 0.1 | | Block x Diet | F(1,15) = 3.2 | p = 0.09 | $\eta_p^2 = 0.2$ | | | |
| 2I&K | SD Intake (1h) | 2-way ANOVA | Diet | F(1,29) = 1.5 | p = 0.2 | Unpaired t-test (Males) | | t(14) = 0.9 | p = 0.4 | d = 0.5 | - | - | SD n = 7/8<br>vHFD n = 9/9 |
| | | | Sex | F(1,29) = 1.3 | p = 0.3 | | | | | | | | |
| | | | Diet x Sex | F(1,29) = 0.3 | p = 0.6 | Unpaired t-test (Females) | | t(15) = 0.8 | p = 0.4 | d = 0.4 | - | - | |
| | SD Intake (2h) | 2-way ANOVA | Diet | F(1,29) = 0.3 | p = 0.6 | Unpaired t-test (Males) | | t(14) = 0.4 | p = 0.7 | d = 0.2 | - | - | |
| | | | Sex | F(1,29) = 0.4 | p = 0.6 | | | | | | | | |
| | | | Diet x Sex | F(1,29) = 0.01 | p = 0.9 | Unpaired t-test (Females) | | t(15) = 0.5 | p = 0.6 | d = 0.2 | - | - | |
| | SD Intake (3h) | 2-way ANOVA | Diet | F(1,29) = 0.09 | p = 0.8 | Unpaired t-test (Males) | | t(14) = 0.6 | p = 0.6 | d = 0.3 | - | - | |
| | | | Sex | F(1,29) = 0.09 | p = 0.8 | | | | | | | | |
| | | | Diet x Sex | F(1,29) = 0.5 | p = 0.5 | Unpaired t-test (Females) | | t(15) = 0.6 | p = 0.6 | d = 0.3 | - | - | |
| | SD Intake (6h) | 2-way ANOVA | Diet | F(1,29) = 1.1 | p = 0.3 | Unpaired t-test (Males) | | t(14) = 0.4 | p = 0.7 | d = 0.2 | - | - | |
| | | | Sex | F(1,29) = 5.1 | p = 0.03 | | | | | | | | |
| | | | Diet x Sex | F(1,29) = 3.0 | p = 0.09 | Unpaired t-test (Females) | | t(15) = 3.3 | p = 0.005 | d = 1.6 | - | - | |
| 2J&L | SD intake rate | 3-way ANOVA | Diet | F(1,29) = 1.1 | p = 0.3 | 2-way ANOVA (Males) | Diet | F(1,14) = 0.1 | p = 0.7 | $\eta_p^2 = 0.6$ | - | - | |
| | | | Sex | F(1,29) = 5.1 | p = 0.03 | | Block | F(1,14) = 23.7 | p < 0.001 | $\eta_p^2 = 0.6$ | | | |
| | | | Block | F(1,29) = 53.3 | p < 0.001 | | Block x Diet | F(1, 14) = 0.5 | p = 0.5 | $\eta_p^2 = 0.03$ | | | |
| | | | Block x Sex | F(1,29) = 1.8 | p = 0.2 | | | | | | | | |
| | | | Block x Diet | F(1,29) = 1.8 | p = 0.2 | 2-way ANOVA (Females) | Diet | F(1,15) = 10.9 | p = 0.005 | $\eta_p^2 = 0.4$ | - | - | |
| | | | Diet x Sex | F(1,29) = 3.0 | p = 0.09 | | Block | F(1,15) = 35.7 | p < 0.001 | $\eta_p^2 = 0.7$ | | | |
| | | | Block x Sex x Diet | F(1,29) = 0.02 | p = 0.9 | | Block x Diet | F(1,15) = 0.2 | p = 0.1 | $\eta_p^2 = 0.1$ | | | |

### Figure 3

| Figure | Data | Test | Factor | F-value or t-value | p-value | Test | Factor | F-value or t-value | p-value | Effect size | Pairwise comparisons | | n (M/F) |
|---|---|---|---|---|---|---|---|---|---|---|---|---|---|
| 3C&E | vHFD Intake (1h) | 2-way ANOVA | Diet | F(1,29) = 0.003 | p = 1.0 | Unpaired t-test (Males) | | t(14) = 0.7 | p = 0.5 | d = 0.4 | - | - | SD n = 7/8<br>vHFD n = 9/9 |
| | | | Sex | F(1,29) = 3.2 | p = 0.08 | | | | | | | | |
| | | | Diet x Sex | F(1,29) = 1.1 | p = 0.3 | Unpaired t-test (Females) | | t(15) = 0.8 | p = 0.4 | d = 0.4 | - | - | |
| | vHFD Intake (2h) | 2-way ANOVA | Diet | F(1,29) = 0.7 | p = 0.4 | Unpaired t-test (Males) | | t(14) = 0.7 | p = 0.5 | d = 0.4 | - | - | |
| | | | Sex | F(1,29) = 1.9 | p = 0.2 | | | | | | | | |
| | | | Diet x Sex | F(1,29) = 0.1 | p = 0.7 | Unpaired t-test (Females) | | t(15) = 0.4 | p = 0.7 | d = 0.2 | - | - | |
| | vHFD Intake (3h) | 2-way ANOVA | Diet | F(1,29) = 2.1 | p = 0.2 | Unpaired t-test (Males) | | t(14) = 0.7 | p = 0.5 | d = 0.4 | - | - | |
| | | | Sex | F(1,29) = 0.1 | p = 0.7 | | | | | | | | |
| | | | Diet x Sex | F(1,29) = 0.08 | p = 0.8 | Unpaired t-test (Females) | | t(15) = 1.4 | p = 0.2 | d = 0.7 | - | - | |
| | vHFD Intake (6h) | 2-way ANOVA | Diet | F(1,29) = 13.6 | p < 0.001 | Unpaired t-test (Males) | | t(14) = 2.5 | p = 0.01 | d = 1.3 | - | - | |
| | | | Sex | F(1,29) = 0.7 | p = 0.4 | | | | | | | | |
| | | | Diet x Sex | F(1,29) = 0.2 | p = 0.7 | Unpaired t-test (Females) | | t(15) = 2.8 | p = 0.01 | d = 1.4 | - | - | |
| 3D&F | vHFD intake rate | 3-way ANOVA | Diet | F(1,29) = 13.6 | p < 0.001 | 2-way ANOVA (Males) | Diet | F(1,14) = 6.2 | p = 0.03 | $\eta_p^2 = 0.3$ | - | - | |
| | | | Sex | F(1,29) = 0.7 | p = 0.4 | | Block | F(1,14) = 54.5 | p = < 0.001 | $\eta_p^2 = 0.8$ | | | |
| | | | Block | F(1,29) = 13.6 | p < 0.001 | | Block x Diet | F(1, 14) = 1.4 | p = 0.3 | $\eta_p^2 = 0.09$ | | | |
| | | | Block x Sex | F(1,29) = 0.02 | p = 0.9 | | | | | | | | |
| | | | Block x Diet | F(1,29) = 1.0 | p = 0.3 | 2-way ANOVA (Females) | Diet | F(1,15) = 7.9 | p = 0.01 | $\eta_p^2 = 0.3$ | - | - | |
| | | | Diet x Sex | F(1,29) = 0.2 | p = 0.7 | | Block | F(1,15) = 87.3 | p < 0.001 | $\eta_p^2 = 0.9$ | | | |
| | | | Block x Sex x Diet | F(1,29) = 0.7 | p = 0.4 | | Block x Diet | F(1, 15) = 0.02 | p = 0.9 | $\eta_p^2 = 0.001$ | | | |
| 3I&K | HFD Intake (1h) | 2-way ANOVA | Diet | F(1,29) = 0.2 | p = 0.7 | Unpaired t-test (Males) | | t(14) = 1.1 | p = 0.3 | d = 0.6 | - | - | SD n = 7/8<br>vHFD n = 9/9 |
| | | | Sex | F(1,29) = 3.1 | p = 0.09 | | | | | | | | |
| | | | Diet x Sex | F(1,29) = 1.6 | p = 0.2 | Unpaired t-test (Females) | | t(15) = 0.6 | p = 0.5 | d = 0.3 | - | - | |
| | HFD Intake (2h) | 2-way ANOVA | Diet | F(1,29) = 0.05 | p = 0.8 | Unpaired t-test (Males) | | t(14) = 1.6 | p = 0.2 | d = 0.8 | - | - | |
| | | | Sex | F(1,29) = 2.4 | p = 0.1 | | | | | | | | |
| | | | Diet x Sex | F(1,29) = 4.6 | p = 0.04 | Unpaired t-test (Females) | | t(15) = 1.5 | p = 0.2 | d = 0.7 | - | - | |
| | HFD Intake (3h) | 2-way ANOVA | Diet | F(1,29) = 0.06 | p = 0.8 | Unpaired t-test (Males) | | t(14) = 1.9 | p = 0.08 | d = 0.9 | - | - | |
| | | | Sex | F(1,29) = 0.5 | p = 0.5 | | | | | | | | |
| | | | Diet x Sex | F(1,29) = 6.8 | p = 0.01 | Unpaired t-test (Females) | | t(15) = 1.8 | p = 0.09 | d = 0.9 | - | - | |
| | HFD Intake (6h) | 2-way ANOVA | Diet | F(1,29) = 2.0 | p = 0.2 | Unpaired t-test (Males) | | t(14) = 2.4 | p = 0.03 | d = 1.2 | - | - | |
| | | | Sex | F(1,29) = 0.01 | p = 0.9 | | | | | | | | |
| | | | Diet x Sex | F(1,29) = 4.3 | p = 0.04 | Unpaired t-test (Females) | | t(15) = 0.5 | p = 0.6 | d = 0.2 | - | - | |
| 3J&L | HFD intake rate | 3-way ANOVA | Diet | F(1,29) = 2.0 | p = 0.2 | 2-way ANOVA (Males) | Diet | F(1,14) = 5.9 | p = 0.03 | $\eta_p^2 = 0.3$ | - | - | |
| | | | Sex | F(1,29) = 0.01 | p = 0.9 | | Block | F(1,14) = 10.6 | p = 0.006 | $\eta_p^2 = 0.4$ | | | |
| | | | Block | F(1,29) = 20.2 | p < 0.001 | | Block x Diet | F(1, 14) = 0.5 | p = 0.5 | $\eta_p^2 = 0.03$ | | | |
| | | | Block x Sex | F(1,29) = 1.1 | p =0.3 | | | | | | | | |
| | | | Block x Diet | F(1,29) = 0.7 | p = 0.4 | 2-way ANOVA (Females) | Diet | F(1,15) = 0.2 | p = 0.7 | $\eta_p^2 = 0.01$ | *Post hoc* Bonferroni (Diet) | 0-3h: p = 0.09<br>3-6h: p = 0.09 | |
| | | | Diet x Sex | F(1,29) = 4.3 | p = 0.04 | | Block | F(1,15) = 9.8 | p = 0.007 | $\eta_p^2 = 0.4$ | | | |
| | | | Block x Sex x Diet | F(1,29) = 4.0 | p = 0.05 | | Block x Diet | F(1, 15) = 6.7 | p = 0.02 | $\eta_p^2 = 0.3$ | | | |

| Figure 4 | | | | | | | | | | | | | |
|---|---|---|---|---|---|---|---|---|---|---|---|---|---|
| **Figure** | **Data** | **Test** | **Factor** | **F-value or t-value** | **p-value** | **Test** | **Factor** | **F-value or t-value** | **p-value** | **Effect size** | **Pairwise comparisons** | | **n (M/F)** |
| **4A** | **Saccharin preference** | 3-way ANOVA | Diet | F(1, 29) = 0.5 | p = 0.5 | 2-way ANOVA (Males) | Diet | F(1, 14) = 0.2 | p = 0.7 | $\eta_p^2$ = 0.01 | | | |
| | | | Sex | F(1, 29) = 1.9 | p = 0.2 | | Concentration | F(1.54, 21.58) = 57.2 | p < 0.001 | $\eta_p^2$ = 0.8 | - | - | |
| | | | Concentration | F(1.84, 53.4) = 125.5 | p < 0.001 | | Diet x Concentration | F(1.54, 21.58) = 0.6 | p = 0.5 | $\eta_p^2$ = 0.04 | | | |
| | | | Diet x Sex | F(1, 29) = 0.05 | p = 0.8 | | | | | | | | |
| | | | Diet x Concentration | F(1.84, 53.4) = 1.0 | p = 0.4 | 2-way ANOVA (Females) | Diet | F(1, 15) = 0.4 | p = 0.6 | $\eta_p^2$ = 0.02 | | | |
| | | | Sex x Concentration | F(1.84, 53.4) = 1.2 | p = 0.3 | | Concentration | F(2.13, 31.88) = 71.1 | p < 0.001 | $\eta_p^2$ = 0.8 | - | - | |
| | | | Diet x Sex x Concentration | F(1.84, 53.4) = 0.8 | p = 0.5 | | Diet x Concentration | F(2.13, 31.88) = 1.3 | p = 0.3 | $\eta_p^2$ = 0.08 | | | |
| **4B** | **Saccharin intake** | 3-way ANOVA | Diet | F(1, 29) = 0.2 | p = 0.6 | 2-way ANOVA (Males) | Diet | F(1, 14) = 0.4 | p = 0.6 | $\eta_p^2$ = 0.03 | | | |
| | | | Sex | F(1, 29) = 1.1 | p = 0.3 | | Concentration | F(1.69, 23.59) = 41.1 | p < 0.001 | $\eta_p^2$ = 0.8 | - | - | |
| | | | Concentration | F(2.11, 61.25) = 95.7 | p < 0.001 | | Diet x Concentration | F(1.69, 23.59) = 0.5 | p = 0.6 | $\eta_p^2$ = 0.04 | | | |
| | | | Diet x Sex | F(1, 29) = 0.2 | p = 0.7 | | | | | | | | SD n = 7/8 vHFD n = 9/9 |
| | | | Diet x Concentration | F(2.11, 61.25) = 0.9 | p = 0.4 | 2-way ANOVA (Females) | Diet | F(1, 15) = 0.005 | p = 1.0 | $\eta_p^2$ = 0 | | | |
| | | | Sex x Concentration | F(2.11, 61.25) = 0.8 | p = 0.5 | | Concentration | F(3, 45) = 56.5 | p < 0.001 | $\eta_p^2$ = 0.8 | - | - | |
| | | | Diet x Sex x Concentration | F(2.11, 61.25) = 1.9 | p = 0.2 | | Diet x Concentration | F(3, 45) = 2.5 | p = 0.07 | $\eta_p^2$ = 0.1 | | | |
| **4C** | **Water intake** | 3-way ANOVA | Diet | F(1, 29) = 0.1 | p = 0.7 | 2-way ANOVA (Males) | Diet | F(1, 14) = 0.01 | p = 0.9 | $\eta_p^2$ = 0.001 | | | |
| | | | Sex | F(1, 29) = 2.4 | p = 0.1 | | Concentration | F(1.36, 19.06) = 45.8 | p < 0.001 | $\eta_p^2$ = 0.8 | - | - | |
| | | | Concentration | F(1.54, 44.77) = 76.6 | p < 0.001 | | Diet x Concentration | F(1.36, 19.06) = 0.4 | p = 0.6 | $\eta_p^2$ = 0.2 | | | |
| | | | Diet x Sex | F(1, 29) = 0.05 | p = 0.8 | | | | | | | | |
| | | | Diet x Concentration | F(1.54, 44.77) = 2.4 | p = 0.1 | 2-way ANOVA (Females) | Diet | F(1, 15) = 0.2 | p = 0.7 | $\eta_p^2$ = 0.01 | | | |
| | | | Sex x Concentration | F(1.54, 44.77) = 2.3 | p = 0.1 | | Concentration | F(1.74, 26.15) = 30.4 | p < 0.001 | $\eta_p^2$ = 0.7 | - | - | |
| | | | Diet x Sex x Concentration | F(1.54, 44.77) = 0.5 | p = 0.6 | | Diet x Concentration | F(1.74, 26.15) = 2.8 | p = 0.09 | $\eta_p^2$ = 0.2 | | | |
| **4D** | **Open field** | 2-way ANOVA Distance | Diet | F(1, 28) = 0.6 | p = 0.5 | Unpaired t-test (Males) | | t(14) = 1.4 | p = 0.2 | d = 0.7 | | | SD n = 7/8 vHFD n = 9/8 |
| | | | Sex | F(1, 28) = 11.5 | p = 0.002 | | | | | | | | |
| | | | Diet x Sex | F(1, 28) = 1.7 | p = 0.2 | Unpaired t-test (Females) | | t(14) = 0.4 | p = 0.7 | d = 0.2 | - | - | |
| | | 2-way ANOVA Entries center | Diet | F(1, 28) = 1.3 | p = 0.3 | Unpaired t-test (Males) | | t(14) = 0.5 | p = 0.6 | d = 0.3 | - | - | |
| | | | Sex | F(1, 28) = 7.6 | p = 0.01 | | | | | | | | |
| | | | Diet x Sex | F(1, 28) = 0.2 | p = 0.7 | Unpaired t-test (Females) | | t(14) = 1.6 | p = 0.1 | d = 0.8 | - | - | |
| | | 2-way ANOVA % time centre | Diet | F(1, 28) = 34.6 | p < 0.001 | Unpaired t-test (Males) | | t(14) = 6.2 | p < 0.001 | d = 3.1 | - | - | |
| | | | Sex | F(1, 28) = 0.1 | p = 0.8 | | | | | | | | |
| | | | Diet x Sex | F(1, 28) = 0.1 | p = 0.8 | Unpaired t-test (Females) | | t(14) = 3.3 | p = 0.006 | d = 1.6 | - | - | |
| **4E** | **Zero maze** | 2-way ANOVA Distance | Diet | F(1, 29) = 0.5 | p = 0.5 | Unpaired t-test (Males) | | t(14) = 1.3 | p = 0.2 | d = 0.6 | - | - | SD n = 7/8 vHFD n = 9/8 |
| | | | Sex | F(1, 29) = 3.5 | p = 0.07 | | | | | | | | |
| | | | Diet x Sex | F(1, 29) = 0.2 | p = 0.6 | Unpaired t-test (Females) | | t(15) = 0.1 | p = 0.9 | d = 0.06 | - | - | |
| | | 2-way ANOVA Entries center | Diet | F(1, 29) = 0.1 | p = 0.8 | Unpaired t-test (Males) | | t(14) = 0.2 | p = 0.9 | d = 0.08 | - | - | |
| | | | Sex | F(1, 29) = 7.8 | p = 0.009 | | | | | | | | |
| | | | Diet x Sex | F(1, 29) = 0.2 | p = 0.7 | Unpaired t-test (Females) | | t(15) = 0.4 | p = 0.7 | d = 0.2 | - | - | |
| | | 2-way ANOVA % time centre | Diet | F(1, 29) = 0.8 | p = 0.4 | Unpaired t-test (Males) | | t(14) = 0.2 | p = 0.9 | d = 0.1 | - | - | |
| | | | Sex | F(1, 29) = 3.8 | p = 0.06 | | | | | | | | |
| | | | Diet x Sex | F(1, 29) = 0.4 | p = 0.5 | Unpaired t-test (Females) | | t(15) = 1.1 | p = 0.3 | d = 0.5 | - | - | |
